# Towards Multimodal Longitudinal Analysis for Predicting Cognitive Decline

**DOI:** 10.1101/2024.11.18.624226

**Authors:** Rayan M. Ashish, Jessica A. Turner

## Abstract

Understanding and predicting cognitive decline in Alzheimer’s disease (AD) is crucial for timely intervention and management. While neuroimaging biomarkers and clinical assessments are valuable individually, their combined predictive power and interaction with demographic and cognitive variables remain underexplored. This study lays the groundwork for comprehensive longitudinal analyses by integrating neuroimaging markers and clinical data to predict cognitive changes over time. Using data from the Alzheimer’s Disease Neuroimaging Initiative (ADNI), we applied feature-driven supervised machine learning techniques for assessing cognitive decline predictability. We hypothesize that combining neuroimaging biomarkers with demographic and clinical assessment variables significantly improves the prediction of cognitive decline in Alzheimer’s disease. Our results show that while imaging biomarkers alone offer moderate predictive capabilities, including key clinical assessment and demographic variables in conjunction with imaging biomarkers significantly improves the model performance. Furthermore, our results indicate that non-imaging variables *alone* can serve as effective and cost-efficient predictors of cognitive decline. This study underscores the need for integrating multi-dimensional data in future longitudinal research to capture time-dependent patterns in cognitive decline and guide the development of targeted intervention strategies. We also introduce NeuroLAMA - an open and extensible data engineering and machine-learning system, to support the continued investigation by the community

## INTRODUCTION

Alzheimer’s disease (AD) is one of the most prevalent neurodegenerative disorders, characterized by progressive cognitive decline and structural brain changes (1). Early detection and accurate monitoring of cognitive impairment are vital for managing disease progression and evaluating the efficacy of therapeutic interventions. Biomarkers, derived from neuroimaging, refer to measurable indicators such as changes in brain volume, activity patterns, or structural and functional connectivity, identified through imaging techniques like Magnetic-Resonance-Imaging (MRI) and Positron-Emission-Tomography (PET). These imaging-based biomarkers provide quantitative measures that reflect underlying neurodegenerative changes, making them invaluable tools for research and clinical practice. Cognitive and functional performance is often assessed using standardized neuropsychological tests, which, when combined with imaging-based biomarkers, offer a comprehensive approach to understanding disease progression and treatment efficacy. The change in cognitive and functional performance over time serves as a measure of disease progression, offering insights into how patients’ cognitive and functional abilities evolve (3). Despite the known value of neuroimaging biomarkers and clinical assessments individually, understanding their predictive relationship and how they interact with other factors such as demographic and cognitive variables remains an area ripe for exploration. Recent studies underscore the importance of integrating neuroimaging biomarkers with clinical assessments and demographic factors to enhance the prediction of cognitive decline. For instance, a 2024 study demonstrated that combining plasma biomarkers with traditional imaging techniques improves the prediction of cognitive decline in non-demented individuals, highlighting the value of a multifaceted approach (4). Similarly, research published in 2023 found that multi-modal neuroimaging signatures, when combined with clinical data, effectively predict cognitive decline in individuals with multiple sclerosis, emphasizing the need to consider various factors in predictive modeling (5). These findings suggest that while neuroimaging and clinical assessments are valuable individually, their predictive power is significantly enhanced when analyzed in conjunction with demographic and cognitive variables.

The source of our data, the Alzheimer’s Disease Neuroimaging Initiative (ADNI) is a longitudinal, multi-center study launched in 2004 with the aim of developing clinical, imaging, genetic, and biochemical biomarkers for the early detection and tracking of Alzheimer’s disease (AD) (6). The study collects data from a wide range of participants, including cognitively normal older adults, individuals with Mild-Cognitive-Impairment (MCI), and patients with AD, enabling researchers to investigate patterns of cognitive decline and identify potential early indicators of the disease. The comprehensive data provided by ADNI includes neuroimaging scans (such as MRI and PET), cognitive assessments, genetic information, and clinical evaluations.

This study investigates the predictive power of different combinations of clinical assessment and demographic variables for changes in cognitive performance. These variables are derived from factors such as the subject’s age, length of formal education, quantitative memory assessment scores, and physical activity levels. The analysis examines their effectiveness when used in conjunction with neuroimaging biomarkers. Additionally, we also assessed the predictive power of using *only* clinical assessment and demographic variables for cognitive decline prediction.

Our primary hypothesis in this study is that non-imaging biomarkers, including demographic and clinical assessment data, can serve as significant and cost-efficient predictors of cognitive decline in Alzheimer’s disease including when used alone. The key takeaway is that while imaging biomarkers alone offer moderate predictive capabilities, adding key clinical assessment and demographic factors significantly enhances model performance and further, strongly suggests that cognitive decline can be accurately predicted using only non-imaging-derived factors. This highlights the need for further investigation into a wider space of such non-imaging variables to better understand their predictive capabilities.

## RESULTS

### Biomarker and Dementia Association

We first analyzed the association between the Spatial-Pattern-of-Abnormality-for-Recognition-of-Early-Alzheimer’s-Disease (SPARE_AD) - a reliable biomarker of brain atrophy, and the change in the Clinical-Dementia-Rating-Sum-of-Boxes (CDRSB) - a quantitative measure of functional impairment (2). The change in CDRSB (across consecutive visits) is denoted by CDRSBDIFF. CDRSB is a standardized cognitive and functional assessment tool used to quantify dementia severity across six domains, including memory, orientation, and problem-solving abilities. SPARE_AD is computed from structural MRI scans that quantifies Alzheimer’s-related brain atrophy. It is applicable to both healthy individuals and those with Alzheimer’s disease, with higher SPARE_AD values indicating greater atrophy, particularly in regions such as the medial temporal lobe, and a stronger association with cognitive decline.

We employed simple linear regression, predicting CDRSBDIFF using SPARE_AD as well as the reverse correlation i.e., predicting SPARE_AD from CDRSBDIFF (Figure 1 (A),(B), respectively). The predictive accuracy in both cases is low to moderate, with R^2^ values of 0.19 and 0.21, respectively. However, the relationships are statistically significant (p = 0.015 and p = 0.022), indicating a weak to moderate but meaningful association despite the modest explanatory power of SPARE_AD alone.

**Figure 1:**
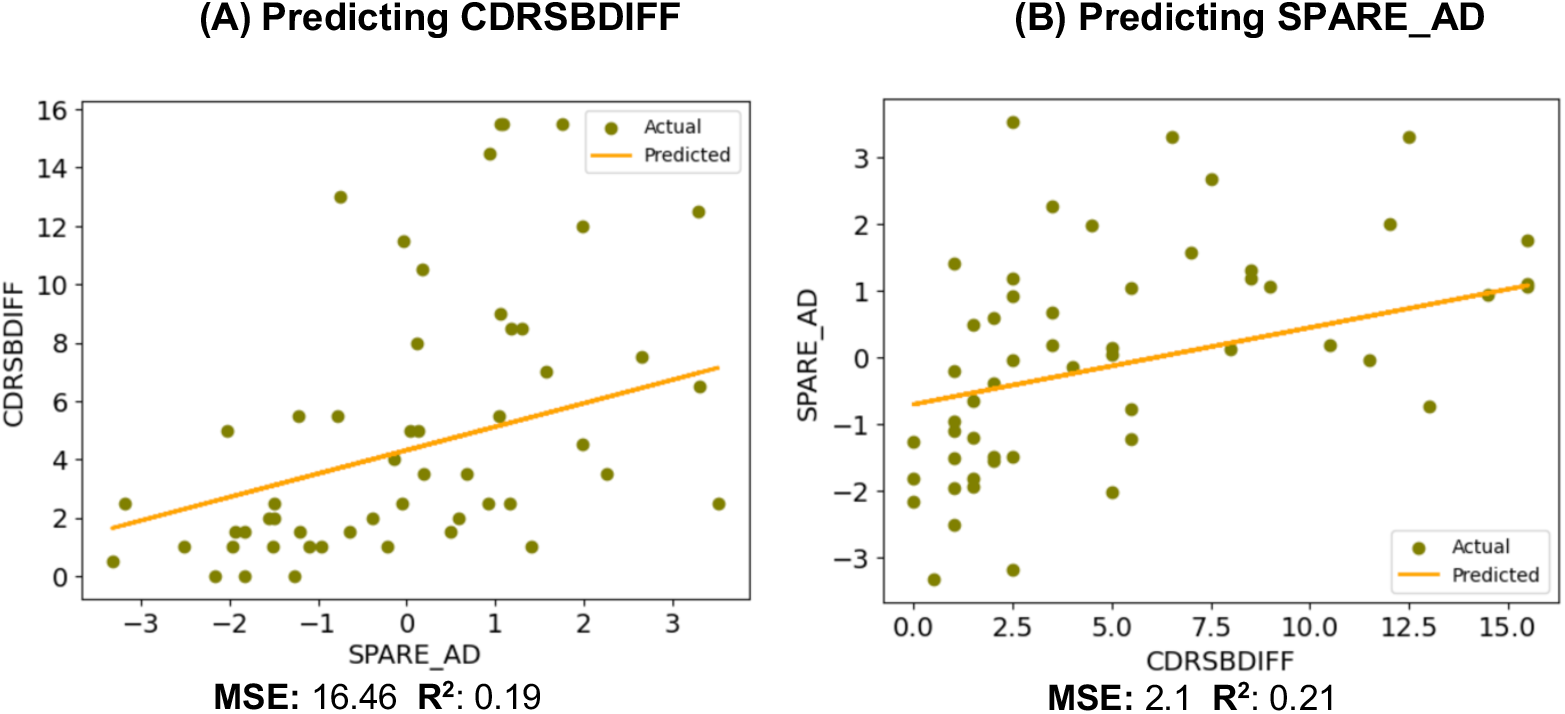
Association of SPARE_AD with CDRSBDIFF.

### Employing Non-imaging Variables in Conjunction

We then evaluated employing additional, non-imaging derived variables in conjunction with SPARE_AD, for predictive cognitive decline, with different sets of variables employed as a group of predictors. These non-imaging variables are the following: AGE_2024, the subject age in 2024; the Mini-Mental-State-Examination Score (MMSCORE), a quantitative measure involving multiple cognitive domains such as orientation, memory etc.; the Abbreviated-Multidimensional-Acculturation-Scale Average (AMASAVG), a self-reported psychological assessment that evaluates cultural adaptation; PTEDUCAT, the number of years of formal education; and PEDAVG, a measure of physical activity levels, derived from self-reported questionnaires. The specific predictor sets that were evaluated are:

1. {SPARE_AD, AGE_2024, PTEDUCAT}
2. {SPARE_AD, MMSCORE, AGE_2024, PTEDUCAT}
3. {SPARE_AD, AMASAVG, MMSCORE, AGE_2024, PTEDUCAT}
4. {SPARE_AD, PEDAVG, MMSCORE, AGE_2024, PTEDUCAT}
5. {PEDAVG, MMSCORE, AGE_2024, PTEDUCAT}

The results are provided for *Random-Forest* regression, which emerged as the best performing model (over Linear-Regression and Decision-Tree). These results include the quality of prediction, as well as the feature importance of individual variables (Figure 2).

**Figure 2:**
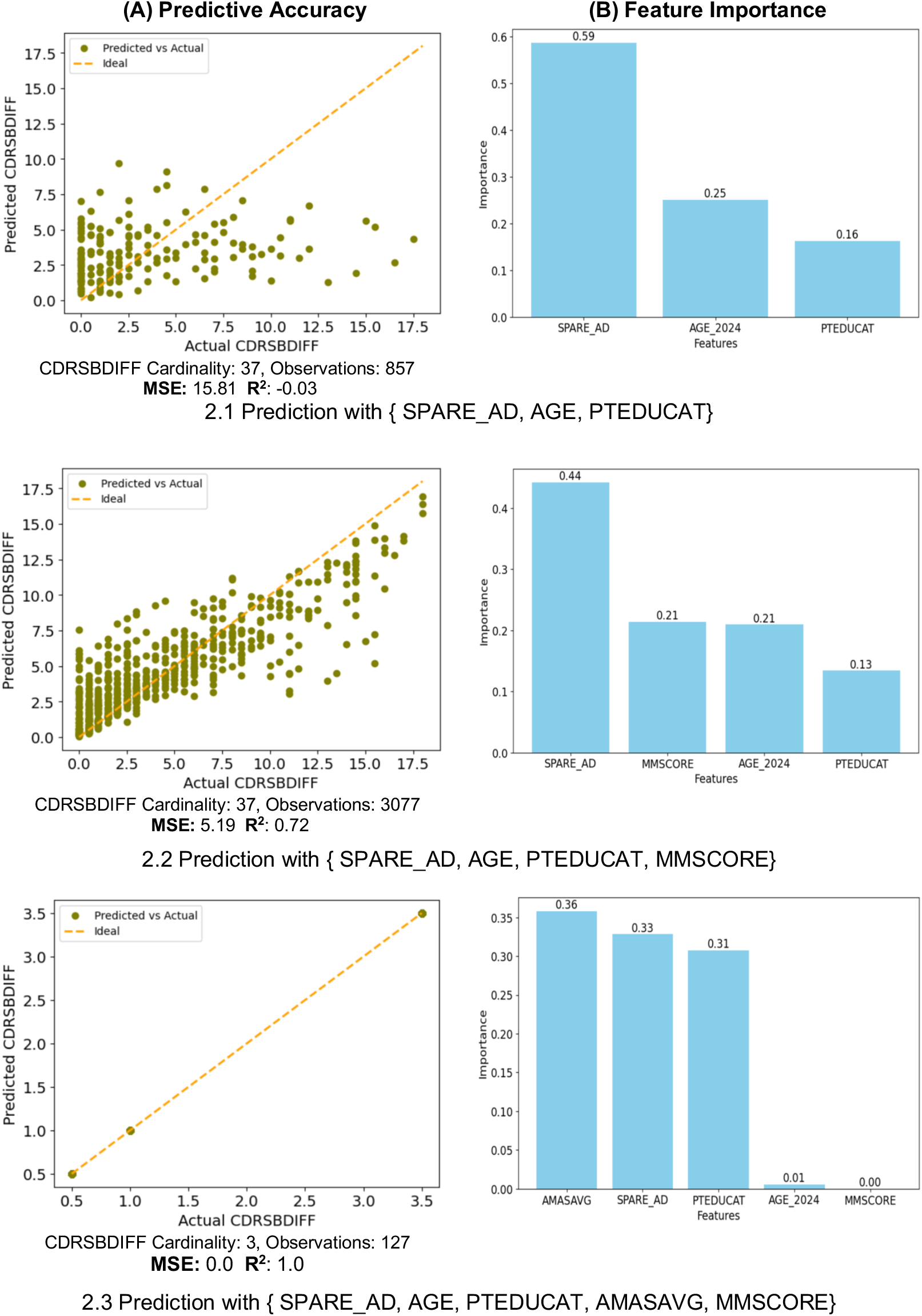
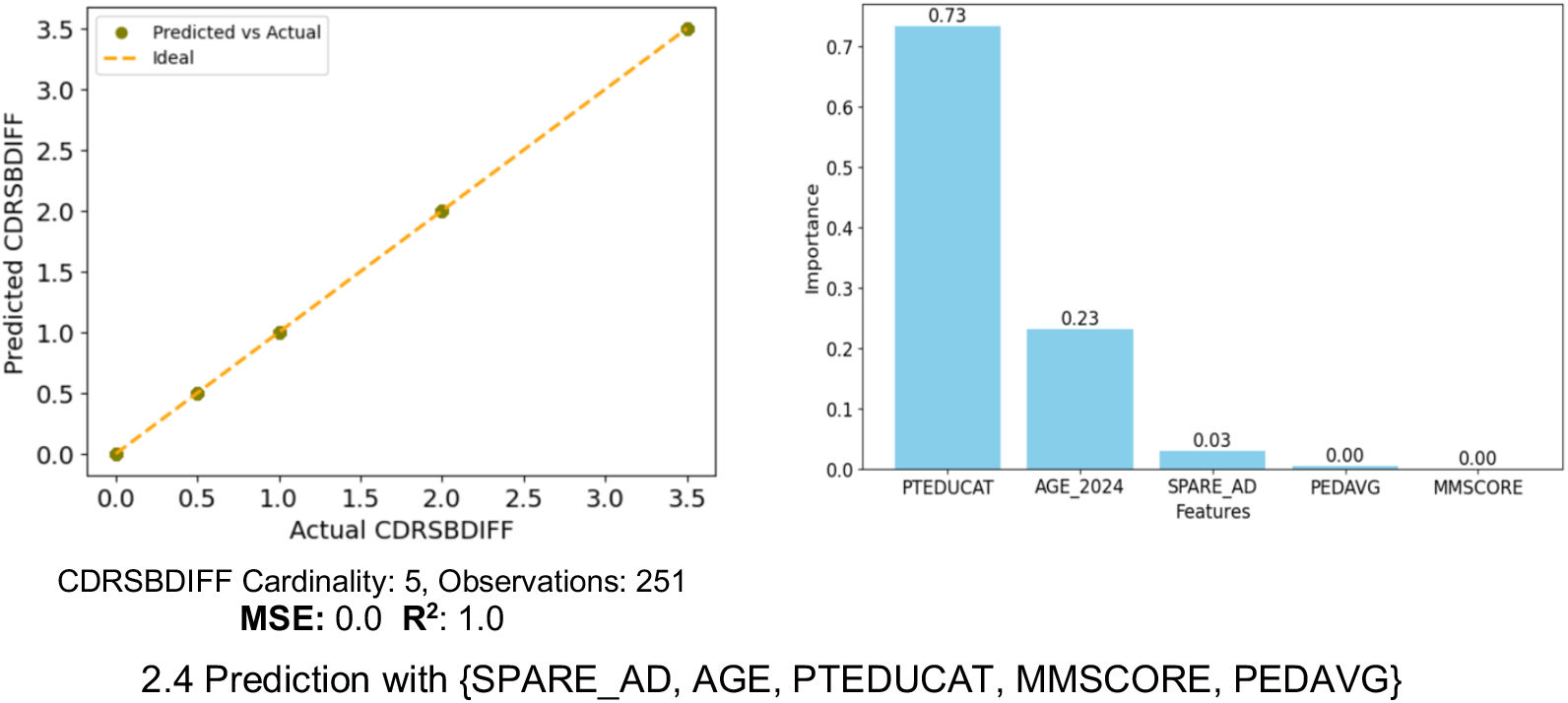
Investigating different variable combinations for predicting CDRSBDIFF.

More than the impact of individual non-imaging variables, the results across predictor sets 1–4 provide valuable insight into how the predictive power of these variables is influenced by their specific combinations. Predictor set 1: {SPARE_AD, AGE_2024, PTEDUCAT} exhibits virtually no predictive power, with an R^2^ of −0.03 and a Mean-Square-Error (MSE) of 15.81 (Figure 2.1(A)). However the addition of the MMSCORE variable in predictor set 2: {SPARE_AD, MMSCORE, AGE_2024, PTEDUCAT}, leads to a significant improvement in prediction accuracy with R^2^ of 0.72 and an MSE of 5.19 (Figure 2.2(A)). Set 3: {SPARE_AD, AGE, PTEDUCAT, AMASAVG, MMSCORE}, with the addition of AMASAVG yields a perfect prediction with an R^2^ of 1.0 and an MSE of 0.0 (Figure 2.3(A)). We must, however, observe this with the context that: a) the observations have been reduced to 127 with this particular combination of variables (versus sets 1 and 2), and b) the cardinality of the target class i.e., CDRSBDIFF has been significantly reduced, from 37 to 3, as well. Finally, set 4: {SPARE_AD, AGE, PTEDUCAT, MMSCORE, PEDAVG} also yields a perfect prediction with an R^2^ of 1.0 and an MSE of 0.0 (Figure 2.4(A)). This is also interpreted with caution with a target class cardinality of 5, as well as a modest sample size of 251.

**Figure 3:**
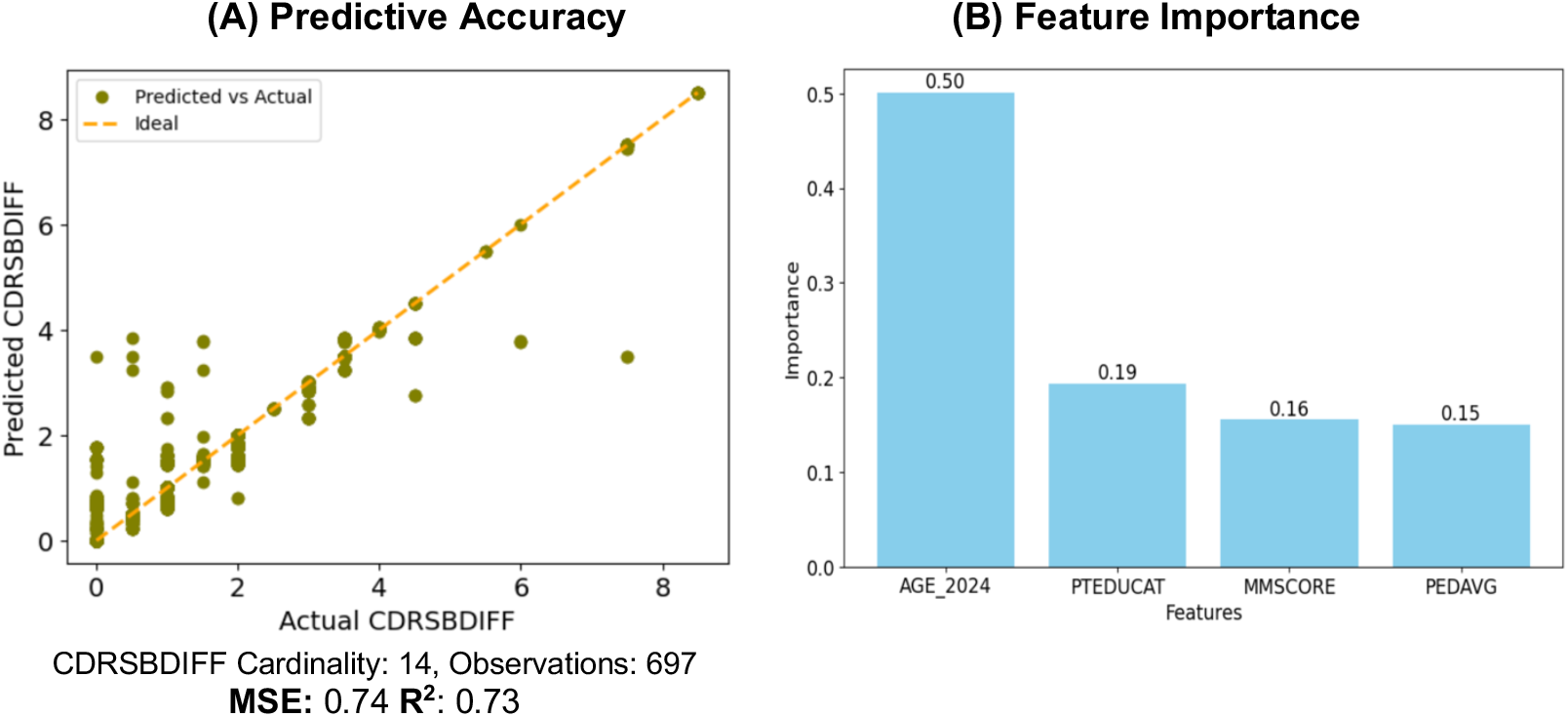
Predicting CDRSBDIFF with non-imaging variables: {AGE, PTEDUCAT, MMSCORE, PEDAVG}

**Figure 4:**
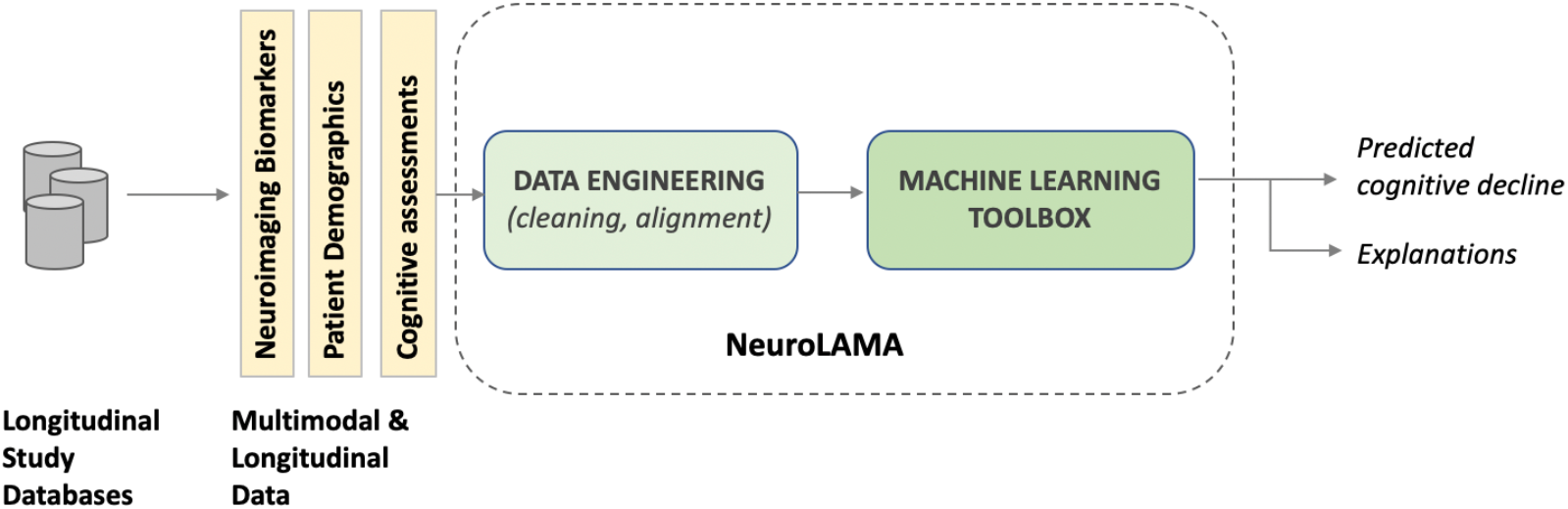
The NeuroLAMA System for Multimodal Longitudinal Predictive Analysis. We utilized the Scikit-Learn framework for the implementation of the various classification based prediction algorithms that we evaluated, of which the best performing ones are reported in the results (14). Our long term goal is that of uncovering specific predictors, including combinations thereof, towards cognitive decline and by leveraging heterogenous potential factors. Our aim with NeuroLAMA is to provide an open machine learning resource to the community, for analysis over prominent datasets of longitudinal Neuroscience studies (such as ADNI).

An interesting observation from the feature importance scores in Predictor Sets 2–4 is that the dominant predictive variable differs across all three cases (Figure 2.2–4(B)). In Set 2, the most influential predictor is SPARE_AD (the imaging variable), whereas in Set 3, it is AMASAVG, and in Set 4, it is PTEDUCAT, both of which are non-imaging variables. However, it is important to note that feature importance values should be interpreted as indicative rather than definitive, as they can be influenced by model parameters, dataset characteristics, and variability in the learning process.

### Employing Non-Imaging Variables Exclusively

We then evaluated variable set 5, which comprises of exclusively non-imaging derived variables. We find the predictive accuracy, with overall R^2^ score of 0.73 and MSE of 0.74, to be high as well as promising (Figure 3(A)). This is taking into account the target class cardinality of 14 and a robust sample size of 697 observations.

### Predicting the Diagnostic Group

Finally, we employed the classifier to predict the *diagnostic group* of the individual i.e., whether the individual is classified as AD/CN/MCI (Alzheimer’s Disease/Cognitively Normal/Mild Cognitive Impairment) based upon the value of CDRSB (2). In this case we employed an *XGBoost* classifier that performed better than Random Forest regression. The results demonstrate a high accuracy for all three groups, with Precision as well as Recall values of 0.90 or above (Table 2). The results for cognitive decline category prediction demonstrate remarkable accuracy, even though only non-imaging variables were used. While this is a preliminary finding based on a limited set of clinical and demographic variables, it suggests that accurate prediction of cognitive decline may be achievable using such readily accessible factors.

## MATERIALS AND METHODS

Our methodology involved the two key components of (i) **Data Engineering:** which is preparing the data from ADNI for our specific machine learning analyses, and (ii) **Machine Learning Implementation:** for conducting the classification based predictive and explainability analyses reported.

We use the CDRSB score from ADNI as a measure of cognitive and functional impairment. CDRSB is a composite score derived from structured clinical interviews assessing multiple domains, including memory, orientation, problem-solving, and daily activities, with higher scores indicating greater impairment (6). To track cognitive decline over time, we compute CDRSBDIFF, which represents the change in CDRSB at each assessment relative to the individual’s first recorded score in the study (baseline). An assessment refers to a clinical cognitive evaluation where CDRSB is recorded. For this analysis, we included individuals with at least six and at most ten assessments over the years, resulting in a cohort of 661 subjects. We also incorporated SPARE_AD, as a neuroimaging biomarker that reflects atrophy patterns in Alzheimer’s disease-vulnerable regions, including the medial temporal lobe, posterior cingulate, and precuneus. The score applies to both healthy controls and individuals with Alzheimer’s disease, with higher values indicating greater similarity to disease-associated brain atrophy. SPARE_AD has been shown to correlate with cognitive decline and disease progression. Table 1 summarizes the age and gender distribution of our cohort and the statistical distribution of CDRSB and CDRSBDIFF in the analyzed data.

**Table 1:** Study cohort. The demographic and cognitive impairment distribution of the ADNI cohort analyzed in this study. CDRSB stands for the Clinical Dementia Rating Sum of Boxes score, and CDRSBDIFF is the difference in the CDRSB score across consecutive visits.

| AGE (years) |  |  | GENDER |  | CDRSB |  |  |  | CDRSBDIFF |  |  |  |
| --- | --- | --- | --- | --- | --- | --- | --- | --- | --- | --- | --- | --- |
| MIN | MAX | AVG | M | F | MIN | MAX | AVG | STD | MIN | MAX | AVG | STD |
| 53 | 109 | 85 | 359 | 302 | 0.0 | 18.0 | 1.88 | 2.75 | 0 | 17.5 | 3.29 | 3.93 |

**Table 2:** Predicting diagnostic group with non-imaging variables. This table provides the results of predicting the *diagnostic group* of an individual with purely non-imaging derived variables. The diagnostic group is determined based on the CDRSB score of the individual with the categories of Cognitively Normal (CN) for *CDRSB* ≤ *0*.*5*, Mild Cognitive Impairment (MCI) for *0*.*5* ≤ *CDRSB* ≤ *4*.*0* and Alzheimer’s Disease (AD) for *CDRSB > 4*.*0*. The prediction accuracy is reported in terms of Precision, Recall, and F1-score for each of the three categories.

| Category | Precision | Recall | F1-score |
| --- | --- | --- | --- |
| AD | 0.90 | 0.92 | 0.91 |
| CN | 0.98 | 0.92 | 0.95 |
| MCI | 0.92 | 0.97 | 0.95 |

### Data Engineering

This involved extracting and integrating data from multiple ADNI tables that included clinical, demographic, neuroimaging, physical activity, and cognitive decline variables. These tables were merged into a single consolidated table with the “RID” (the subject identifier) as the unique key to join records across tables. Finally, and as mentioned earlier, we further filtered this integrated table to only retain subjects who had at least six and at most ten recorded assessments. The data engineering functionality was implemented in Python (12) and using the Pandas data processing library (13).

### Machine Learning Implementation

NeuroLAMA (Neuro Longitudinal and Multimodal Analysis) is an open and extensible machine-learning system that we have designed and developed for such analyses (Figure 4) (15).

## DISCUSSION

The findings from this study highlight the importance of integrating neuroimaging biomarkers, cognitive scores, and demographic variables to predict changes in cognitive function, as measured by CDRSBDIFF. The analyses demonstrate that SPARE_AD, a robust imaging biomarker reflecting Alzheimer’s-related brain atrophy, is limited as a predictor of cognitive decline. However, the predictive power is enhanced when additional variables, such as age, education level, and MMSCORE, are included. This multi-dimensional approach aligns with the understanding that cognitive decline is influenced by an interplay of structural brain changes, cognitive reserve, and demographic factors. Since SPARE_AD alone provides only a partial view of disease progression, incorporating non-imaging variables is essential for improving predictive accuracy. The results with non-imaging variables demonstrate that accurate prediction of cognitive status across AD, MCI, and CN groups is achievable using age, education, cognitive scores, and physical activity data. This finding supports the need for further investigation into a broader range of such accessible variables. If validated, it could significantly impact the development of scalable and resource-efficient screening tools for clinical and community settings.

Despite the promising results, some limitations should be noted. While the models performed well on the provided data, external validation with other cohorts is necessary to confirm the generalizability of the findings. Future studies should incorporate larger and more balanced datasets to mitigate the potential impact of small target class cardinality and limited sample sizes in certain predictor sets, ensuring more robust and reliable predictive performance. Additionally, potential confounding factors not included in the current analysis, such as *lifestyle attributes* (7) as well as *genetic predispositions* (such as APOE-ε4 status) (8), may further elucidate the complex relationships affecting cognitive decline.

Given the longitudinal nature of the data used in this study, future work should emphasize building predictive models that leverage the temporal dimension to enhance understanding of disease progression over time. This essentially implies that the models must be capable of predicting cognitive decline in future time states (years), given the decline trajectory history thus far and the associated demographic variables. Advanced machine learning approaches, such as *Recurrent-Neural-Networks* (RNNs) and *Transformer Models* (9), could be employed to model the *sequential* aspects of cognitive decline and brain changes. These methods would be particularly effective in capturing time-dependent patterns that static models may overlook, offering a more dynamic prediction of future cognitive states. Further research can also investigate the integration of additional longitudinal variables that contribute to cognitive resilience and decline, such as physical activity patterns, social engagement, and comprehensive lifestyle measures. Including genetic information (e.g., APOE-ε4 genotype) and other biomarkers (e.g., tau and amyloid levels) could provide a more detailed picture of the interplay between genetic risk factors and observed brain and cognitive changes over time. Moreover, longitudinal data can be used to analyze how changes in variables like SPARE_AD interact with shifts in *cognitive reserve* (10) and other protective factors over multiple time points. Understanding these interactions will be essential for developing targeted interventions aimed at delaying the onset of cognitive decline and mitigating the impact of Alzheimer’s disease. Finally, from a longitudinal data analysis perspective, we have employed the subject age in year 2024 (versus age at time of each visit). Since all study data comes from ADNI, using age in 2024 provides a uniform reference point across participants, ensuring consistency in comparisons.

On the machine-learning aspect, Random Forest regression performed better than XGBoost in our study. This is likely so because it is more stable and less sensitive to small variations in the data. XGBoost on the other hand builds predictions in a step-by-step manner which can lead to instability. Random Forest takes multiple independent views of the data and combines them, leading to more balanced and reliable predictions, especially when dealing with moderate sample sizes and diverse variables such as in our study.

Our continued work will focus on developing both the machine-learning as well as the clinical analysis thrusts of the work. In machine-learning we are working on developing and providing more advanced analysis algorithms, specifically RNNs and Transformers, in the NeuroLAMA system. In clinical analysis our focus will be on applying these longitudinal models to diverse populations, to ensure findings are applicable across different racial, ethnic, and socio-economic groups. This would enhance the equity and generalizability of the predictive models and inform culturally tailored interventions that address the unique needs of various populations.

## ACKNOWLEDGEMENTS

Data used in preparation of this article were obtained from the Alzheimer’s Disease Neuroimaging Initiative (ADNI) database (adni.loni.usc.edu). As such, the investigators within the ADNI contributed to the design and implementation of ADNI and/or provided data but did not participate in analysis or writing of this report. A complete listing of ADNI investigators can be found at: http://adni.loni.usc.edu/wp-content/uploads/how_to_apply/ADNI_Acknowledgement_List.pdf

